# Integrating omics approaches to discover and prioritize candidate genes involved in oil biosynthesis in soybean

**DOI:** 10.1101/2021.08.05.455283

**Authors:** Dayana K. Turquetti-Moraes, Kanhu C. Moharana, Fabricio Almeida-Silva, Francisnei Pedrosa-Silva, Thiago M. Venancio

**Affiliations:** Laboratório de Química e Função de Proteínas e Peptídeos, Centro de Biociências e Biotecnologia, Universidade Estadual do Norte Fluminense Darcy Ribeiro, Campos dos Goytacazes, RJ, Brazil

**Keywords:** Seed oil content, fatty acids, differentially expressed genes, quantitative trait loci, genome-wide association study

## Abstract

Soybean is one of the major sources of edible protein and oil. Oil content is a quantitative trait that is significantly determined by genetic and environmental factors. Over the past 30 years, a large volume of soybean genetic, genomic, and transcriptomic data have been accumulated. Nevertheless, integrative analyses of such data remain scarce, in spite of their importance for crop improvement. We hypothesized that the co-occurrence of genomic regions for oil-related traits in different studies may reveal more stable regions encompassing important genetic determinants of oil content and quality in soybean. We integrated publicly available data, obtained with distinct techniques, to discover and prioritize candidate genes involved in oil biosynthesis and regulation in soybean. We detected key fatty acid biosynthesis genes (e.g., BCCP and ACCase, FADs, KAS family proteins) and several transcripton factors, which are likely regulators of oil biosynthesis. In addition, we identified new candidates for seed oil accumulation and quality, such as Glyma.03G213300 and Glyma.19G160700, which encode a translocator protein and a histone acetyltransferase, respectively. Further, oil and protein genomic hotspots are strongly associated with breeding and not with domestication, suggesting that soybean domestication prioritized other traits. The genes identified here are promising targets for breeding programs and for the development of soybean lines with increased oil content and quality.

## INTRODUCTION

Soybean (*Glycine max* (L.) Merr.) is a major source of protein and edible oil worldwide, constituting a key factor in human and animal nutrition. With 17% to 22% seed oil content, soybean is also widely used for industrial applications and biodiesel production (Abdelghany et al., 2020). Soybean seed oil consists of triacylglycerol (TAG) ester molecules that accumulate fatty acids (FAs) (Thelen & Ohlrogge, 2002), particularly palmitic (C16:0), stearic (C18:0), oleic (C18:1), linoleic (C18:2), and linolenic (C18:3) acids. The proportion of these FAs typically determines oil quality (Clemente & Cahoon, 2009). For example, polyunsaturated fatty acids (PUFAs) are beneficial for human health (Sacks et al., 2017), although the unsaturation degree and positions determine oil melting point (Voelker & Kinney, 2001). High PUFA levels, particularly that of linolenic acid, increase oil auto-oxidation and reduce its useful life. Hence, a major goal in soybean genetic improvement is to increase oil content and quality (e.g. increasing C18:1 content) (Clemente & Cahoon, 2009; Haun et al., 2014; Liu et al., 2014), including the discovery of important genes involved in such phenotype (Li et al., 2017; Liu et al., 2014; Zhang et al., 2019b; Zhang et al., 2016).

Oil biosynthesis involves different cell compartments and comprises a complex gene network controlled by several quantitative trait loci (QTL) that are influenced by genetic and environmental factors (Bates et al., 2013; Collard & Mackill, 2008; Schmidt & Herman, 2008). In plants, *de novo* FA synthesis within plastids occurs through the coordination of several metabolic pathways including the Calvin cycle, glycolysis, starch metabolism and the pentose phosphate pathway (Bates et al., 2013; Gupta et al., 2017). TAGs are then assembled within the endoplasmic reticulum and stored in oil bodies (Bates et al., 2013; Marchive et al., 2014). Although well characterized in *Arabidopsis*, the genes involved in acyl-lipid metabolism are not fully understood in soybean (Liu et al., 2020; Marchive et al., 2014). The difficulty to functionally characterize these genes in soybean can be partially explained by the high prevalence (i.e., ∼75%) of protein-coding genes in multigene families, mainly because of two whole-genome duplication events (Schmutz et al., 2010).

Over the past decades, several groups have explored the genomic complexity of oil-related traits in soybean through linkage mapping (Akond et al., 2014; Bachlava et al., 2009; Diers et al., 1992; Eskandari et al., 2013; Priolli et al., 2015; Qi et al., 2011; Vaughn et al., 2014) and association mapping (i.e., Genome Wide Association Studies, GWAS) (Leamy et al., 2017; Li et al., 2018; Zhang et al., 2018; Zhou et al., 2015). Detection of consensus QTL have been used to define more stable QTL, i.e., those recurrently found across different environments, often referred to as meta-QTL (MQTL) (Gong et al., 2018; Qin et al., 2018; Van & McHale, 2017). Even though these studies have been extremely important in finding key genes involved in agronomic traits, they often reveal long genomic segments comprising many genes, requiring additional information to pinpoint causative genes or alleles. A rich source of additional data can be found in transcriptomic studies, which have remarkably accumulated over the past few years (Almeida-Silva et al., 2021; Bellieny-Rabelo et al., 2016; Lu et al., 2016b). Recently, our group developed a comprehensive Soybean Expression Atlas with 1,298 RNA-seq samples that can be used to investigate gene expression across different tissues and conditions (Machado et al., 2020). This collection has also been used to build gene co-expression networks (Almeida-Silva et al., 2020), which are instrumental in uncovering important evolutionary trends among duplicated genes.

In spite of the large volume of association and linkage mapping, genomic, and transcriptomic datasets, integrative approaches remain scarce (Liu et al., 2020; Niu et al., 2020; Ronne et al., 2019), resulting in an incomplete picture of metabolic and regulatory genes determining soybean oil content. Here, we integrate large-scale datasets from various sources to define stable genomic regions and identify the most promising genes involved in oil-related traits therein. The integrative strategy implemented here allowed the recovery of genes known to be important for oil synthesis, as well as novel candidate genes to be prioritized in experimental validation studies and in future crop improvement programs.

## RESULTS AND DISCUSSION

### Literature mining for QTL, MQTL, GWAS, selective sweep regions, and genes associated with oil-related traits

A total of 478 QTL controlling oil-related traits (Figure 1A) were retrieved from SoyBase (soybase.org) (Supplementary Figure S1). We performed an initial exploratory analysis of the distribution of these QTL along the soybean genome. The number of QTL per chromosome ranged from 12 (in Gm04 and Gm22) to 42 (in Gm15), with an average of 23.9 QTL per chromosome. The oil QTL sizes range from 0.004 Mb to 47.77 Mb, with an average length of 5.45 Mb. The largest oil QTL (47.77 Mb), comprising more than two thousand genes, was greater than the average size (47.46 Mb) of the soybean chromosomes (Figure 1B; Supplementary table S1).

**Figure 1.**
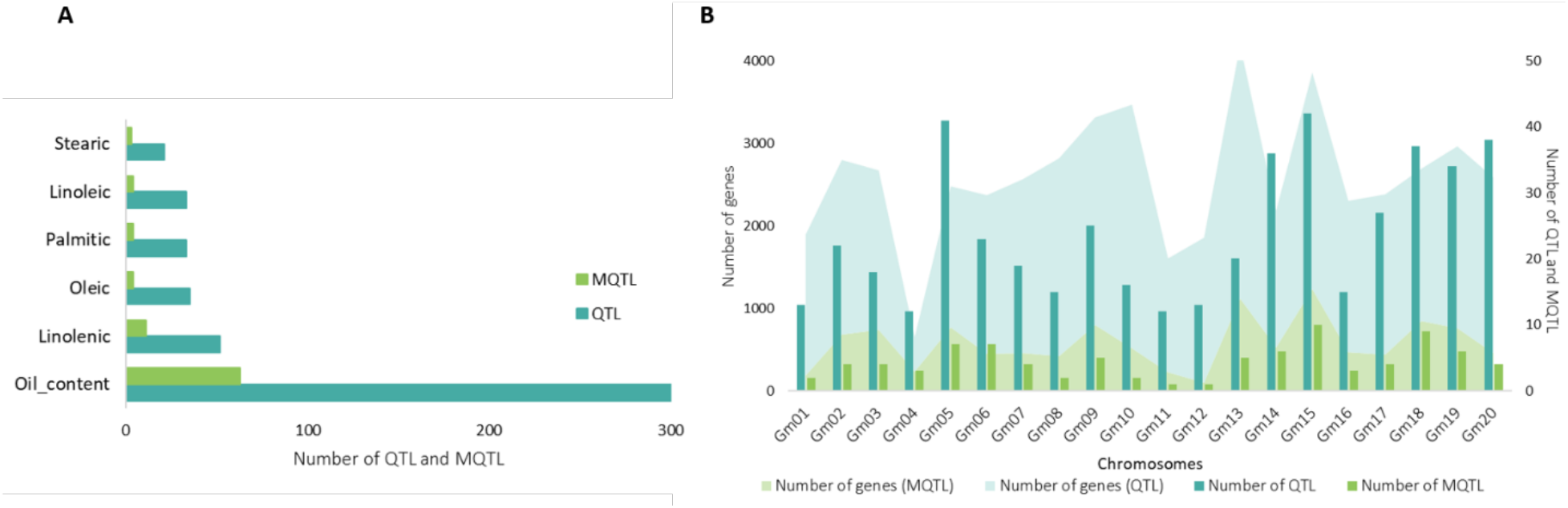
Number of QTL and MQTL for each trait and their distribution on 20 chromosomes of soybean. **A**. Number of QTL and MQTL for oil-related traits. **B**. Distribution of QTL and MQTL per chromosome (bars) and their respective number of genes (shaded regions).

Large confidence intervals are one of the main limitations of QTL analysis, as they make the identification of causal genes a very challenging task (Borevitz & Nordborg, 2003; Collard et al., 2008; Leamy et al., 2017). For example, the 478 QTL collected here encompass 40,268 genes, which correspond to 71.8% (40,268 / 56,044) of the soybean protein-coding genes. Since many oil-related QTL are available, MQTL analyses can be used to better resolve intervals and help identify effective candidate genes (Goffinet et al., 2000). Qi et al. found 89 MQTLs for oil-related traits (Qi et al., 2018). We found that 97.38% (11,104 / 11,403) of the genes in MQTL intervals reported by Qi et al. are also in our QTL database (Figure 1A, B; Supplementary table S1), which is a high correspondence given the several integrated studies, methods, and different genome assembly versions used.

The progress in DNA sequencing technologies significantly improved the identification of single nucleotide polimorfisms (SNPs) in the soybean genome (Song et al., 2013, 2016), paving the way for GWAS (Daware et al., 2017; Mackay et al., 2009) and accelerating the identification of genes with agronomic relevance. We searched the literature and retrieved GWAS data from 15 publications, comprising a total of 458 SNPs significantly associated with oil-related traits (Table 1; Supplementary figure S1). These SNPs allowed us to retrieve 344 statistically significant regions, encompassing 6,804 protein-coding genes (12.14% of the soybean protein-coding genes) (Supplementary table S1 and S2).

**Table 1.**
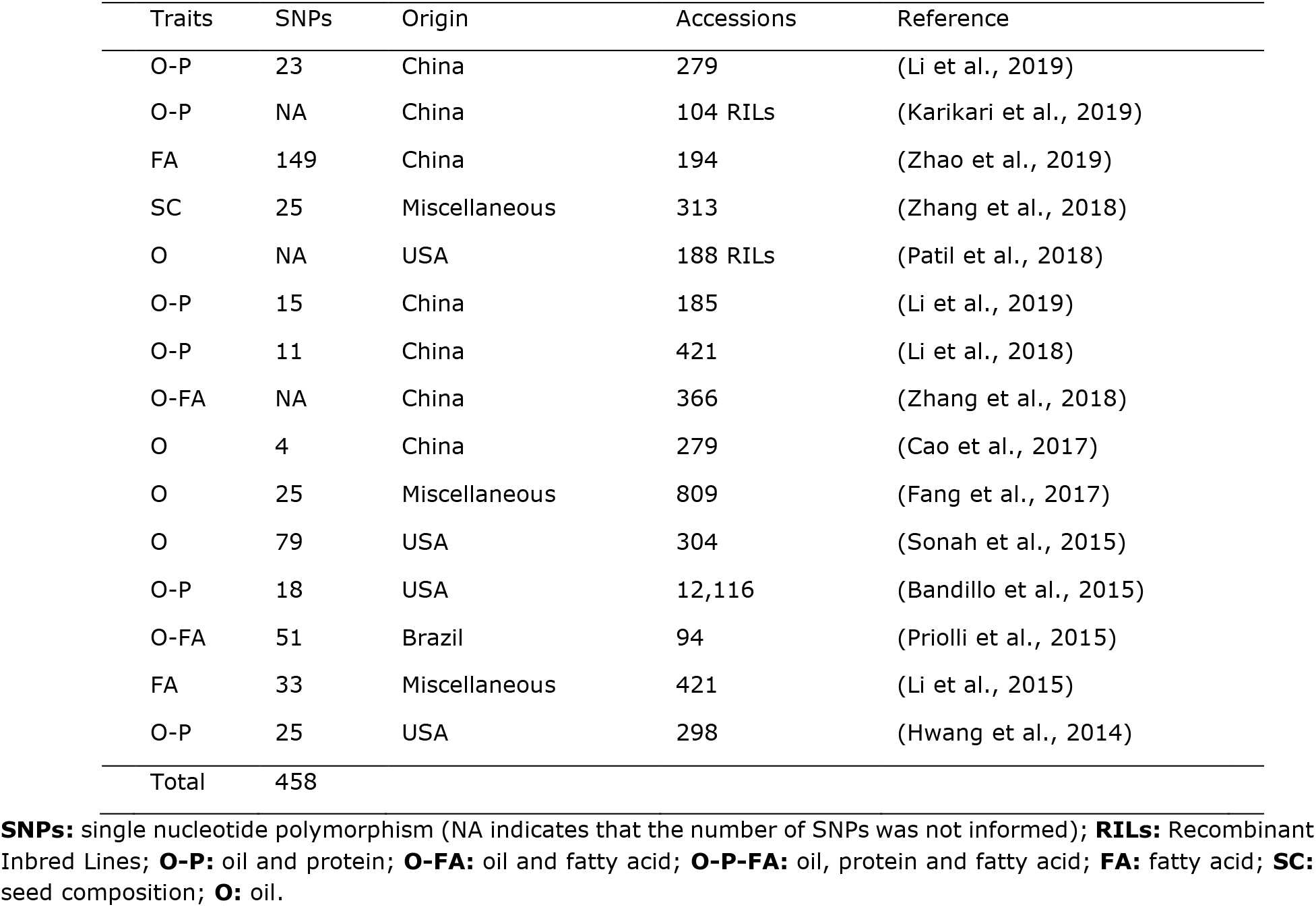
Database of 15 Genome Wide Association Studies (GWAS) used in our analysis

| Traits | SNPs | Origin | Accessions | Reference |
| --- | --- | --- | --- | --- |
| O-P | 23 | China | 279 | (Li et al., 2019) |
| O-P | NA | China | 104 RILs | (Karikari et al., 2019) |
| FA | 149 | China | 194 | (Zhao et al., 2019) |
| SC | 25 | Miscellaneous | 313 | (Zhang et al., 2018) |
| O | NA | USA | 188 RILs | (Patil et al., 2018) |
| O-P | 15 | China | 185 | (Li et al., 2019) |
| O-P | 11 | China | 421 | (Li et al., 2018) |
| O-FA | NA | China | 366 | (Zhang et al., 2018) |
| O | 4 | China | 279 | (Cao et al., 2017) |
| O | 25 | Miscellaneous | 809 | (Fang et al., 2017) |
| O | 79 | USA | 304 | (Sonah et al., 2015) |
| O-P | 18 | USA | 12,116 | (Bandillo et al., 2015) |
| O-FA | 51 | Brazil | 94 | (Priolli et al., 2015) |
| FA | 33 | Miscellaneous | 421 | (Li et al., 2015) |
| O-P | 25 | USA | 298 | (Hwang et al., 2014) |
| Total | 458 |  |  |  |

**SNPs:** single nucleotide polymorphism (NA indicates that the number of SNPs was not informed); **RILs:** Recombinant Inbred Lines; **O-P:** oil and protein; **O-FA:** oil and fatty acid; **O-P-FA:** oil, protein and fatty acid; **FA:** fatty acid; **SC:** seed composition; **O:** oil.

Selective sweep is a process by which a strongly beneficial mutation spreads in a population, increasing the frequency of linked neutral variants in a specific region and dramatically reducing genetic variation in its vicinity (Chen et al., 2010; Stephan, 2019). Allelic variation is lower in domesticated soybean accessions than in its wild relative *Glycine soja*, most likely as a result of strong genetic bottlenecks, such as domestication and selective breeding (Hyten et al., 2006; Liu et al., 2020). It is also clear that oil content was a major target of artificial selection, resulting in increased oil content in cultivated soybean seeds (Wen et al., 2015; Zhou et al., 2015). Hence, domestication and breeding can be used as a model to uncover genes involved in recently selected traits (e.g. high seed oil content) through the identification of selective sweep regions (Chen et al., 2010). Zhou et al. characterized several selective signals related to domestication and breeding through genomic analyses of 302 wild, landrace, and improved soybean accessions (Zhou et al., 2015). We retrieved a total of 2,230 genes within such selective sweep regions, which were categorized as selected during domestication (59.24%), breeding (42.64%) or both (2.60%) (Supplementary table S1; Supplementary figure S2).

We also gathered other datasets to enrich our analyses (Supplementary figure S2): annotated transcription factors (TFs) (Moharana & Venancio, 2020; Niu et al., 2020; Yao et al., 2020); hub genes from co-expression modules for oil-related traits (Qi et al., 2018; Yang et al., 2019); differentially expressed genes (DEGs) between high and low-oil soybean accessions (Niu et al., 2020) and DEGs for a critical period of oil accumulation during soybean seed development (Yang et al., 2019). We leveraged these complementary datasets to better understand the roles of these genes and prioritize candidates for crop improvement strategies, as described below.

### RNA-Seq analyses and the identification of promising candidate genes for oil accumulation and quality

A recent study have identified 126 DEGs, during seed development of a single soybean line with ∼19% of oil content in a critical period of oil accumulation in seeds (Yang et al., 2019). Another study reported 359 DEGs, from comparisons among six contrasting accessions for oil content (11.9 to 12.5%; 17.2 to 17.8% and 20.9 to 22.3%) during seed development (Niu et al., 2020). By comparing the DEGs from both studies, we found only 16 genes in common. Glyma.01G227900 and Glyma.05G013800 encode steroleosin and oleosin, respectively; Glyma.15G105900, Glyma.19G028800, Glyma.20G111000, and Glyma.13G010100 encode a glucose-6-phosphate/phosphate translocator 2 (GPT2), a biotin carboxyl carrier protein (BCCP), a fatty acid desaturase (FAD2-1B), and a long chain acyl-CoA synthesis (LACS8), respectively; Glyma.15G105100 encode an aluminum-induced protein (AILP1); Glyma.20G110900 (an ortholog of AT5G04750) encodes a mitochondrial F1F0-ATP synthase inhibitor factor 1 that has been recently proposed to be crucial for plant growth and responses to abscisic acid (ABA) in *A. thaliana* (Chen et al., 2020); Glyma.08G064400 encodes a protein of unknown function. We also found TFs from the following families: bZIP (Glyma.02G058800 and Glyma.10G071700), NF-YA (Glyma.02G303800), C3H (Glyma.06G290100), GRF (Glyma.07G038400), B3 (Glyma.08G357600), and TALE (Glyma.17G132600). Interestingly, 15 of the 16 genes listed above are located in QTL, GWAS or selective sweep regions (Figure 2). Unexpectedly, only Glyma.13G010100 was found in MQTL regions. This result not only enforces the need for integrating different sources of genomic information, but also indicates that MQTL cannot directly displace QTL data in integrative analyses. Notably, 50%, 43.75% and 6.25% of these 16 genes encode oil biosynthesis or storage genes, TFs and genes of unknown function, respectively (Figure 2; Supplementary table S3). This remarkable representation support TFs as key regulators of transcriptional programs involved in oil accumulation during seed development. Originally, among the DEGs reported by Yang et al. and Niu et al., 9.68% and 29.33% are TFs, respectively (Niu et al., 2020; Yang et al., 2019). This trend has prompted us to further investigate TFs related with oil content in soybean, which are discussed in the next section.

**Figure 2.**
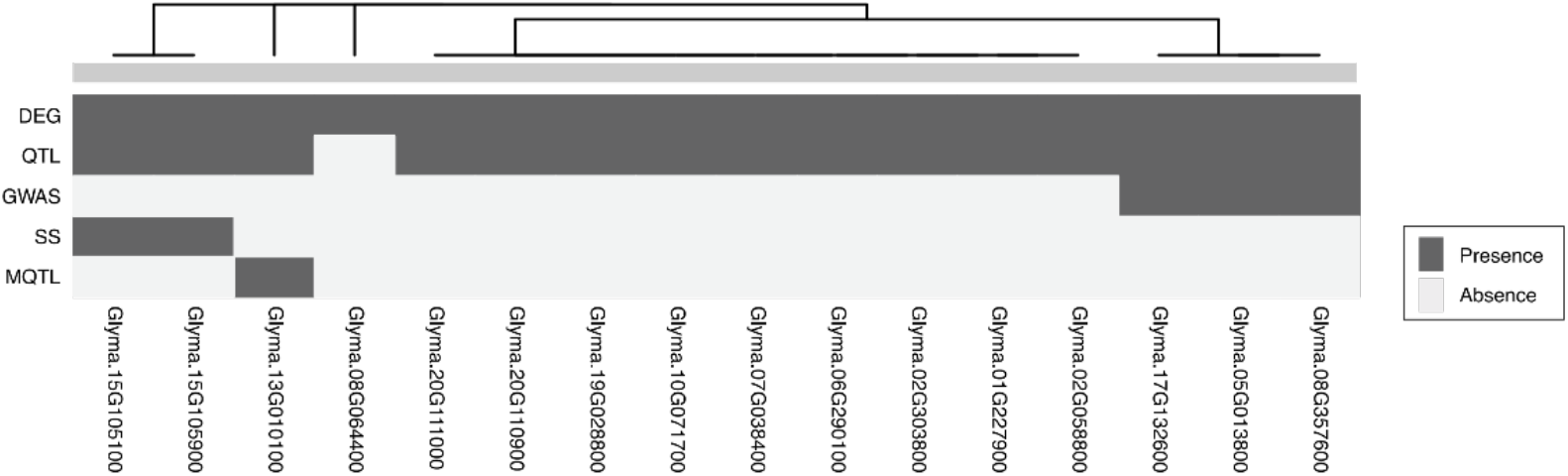
Integration of the genes found by Yang et al. and Niu et al. (Niu et al., 2020; Yang et al., 2019). The heatmap shows the presence/absence of these genes in genomic regions for oil-related traits. DEG, differentially expressed gene. QTL, quantitative trait loci. GWAS, genome wide association studies. SS, selective sweep regions. MQTL, meta-QTL.

Interestingly, 6 of the 16 genes reported above (i.e. GPT2, BCCP, FAD2-1B, LACS8, steroleosin and oleosin) are well described as involved in oil biosynthesis and storage (Haun et al., 2014; Li et al., 2016; Salie et al., 2016a). GPT2 encodes a plastid transporter that imports glucose-6-phosphate into plastids, fueling FA synthesis. GPT2 was likely selected during soybean domestication (Figure 2). GPT2 is 9 times more expressed in oil palm (*Elaeis guineensis*), which accumulates up to 90% oil in its mesocarp, than in date palm (*Phoenix dactylifera*), which stores almost exclusively carbohydrates (Bourgis et al., 2011). BCCP encodes a subunit required for ACCase activity, which catalyzes the committed step of de novo FA synthesis (Salie & Thelen, 2016b). FAD2 encodes a FA desaturase that catalyzes the formation of C18:2 from C18:1 (Haun et al., 2014), while LACS8 is involved in FA export from plastid to TAG synthesis (Li et al., 2016). It has been proposed that the upregulation of LTPs (lipid transporters) and LACs could improve the efficiency of lipid transport and increase oil content (Koo et al., 2004; Manan et al., 2017; Niu et al., 2020). During seed development, there is the deposition of oil bodies, which are TAGs surrounded by a monolayer membrane containing steroleosin and oleosin, among other proteins (Lin et al., 2002; Schmidt et al., 2008). Interestingly, Glyma.20G110900 (related to aerobic respiration) and Glyma.08G064400 (unknown function) have been previously associated to oleic acid content in soybean (Liu et al., 2020; Niu et al., 2020).

We found that Glyma.03G213300 (an ortholog of AT2G47770), which encodes a membrane translocator protein (TSPO), was upregulated in a critical period of oil accumulation in seeds (Yang et al., 2019). Located at the membrane of the endoplasmic reticulum and Golgi complex (Guillaumot et al., 2009), TSPO is expressed in *Arabidopsis* seeds and induced by osmotic treatment, salt stress and ABA (Guillaumot et al., 2009). Further, TSPO is involved in energy homeostasis by promoting the accumulation of FAs and oil bodies in mature seeds (Jurkiewicz et al., 2018). Accordingly, we found that TSPO expression is greater in soybean seed tissues (Supplementary figure S3), leading us to hypothesize it as a strong candidate for oil biosynthesis to be prioritized in experimental validation.

These results corroborate the importance of integrating different datasets to find relevant candidate genes, since genes known to act in oil biosynthesis are often not detected by all methods because of technical limitations (Korte & Farlow, 2013; Mackay et al., 2009) or biological contexts. For example, several classic genes involved in oil biosynthesis were not found in GWAS or MQTL regions (e.g. Glyma.19G028800 and Glyma.20G111000). On the other hand, the integration of transcriptomic data can help discriminating interesting candidates from large QTL intervals (Figure 2; Supplementary table S3).

### Transcription factors in genomic regions for oil-related traits

TFs integrate various signals that coordinate metabolic pathways, including oil biosynthesis (Manan et al., 2017). Despite the knowledge about many TFs involved in oil biosynthesis (Kanai et al., 2019; Pham et al., 2012; Sandhu et al., 2007), several regulators and their regulatory interactions remain unknown (Kong et al., 2020; Wang & Komatsu, 2017). To this end, we investigated the occurrence of soybean TFs in oil-related regions, i.e., QTL, MQTL, GWAS or selective sweep regions. Out of the 3,450 unique TFs considered here, 77.53% were within genomic regions for oil-related traits (Supplementary table S4; Supplementary figure S2). Because of the reasons outlined above, this number is obviously overestimated.

GmZF351 (Glyma.06G290100) increases oil content in soybean seeds by promoting the expression of the WRI1 TFs (Glyma.15G221600; Glyma.08G227700) and other lipid biosynthesis genes, namely BCCP2 (Glyma.19G028800), KASIII (Glyma.15G003100), TAG1 (Glyma.17G053300; Glyma.13G106100) and OLEO2 (Glyma.16G071800; Glyma.19G063400) (Li et al., 2017). GmZF392 (Glyma.12G205700), a homolog of GmZF351, is also important for lipid accumulation. GmZF392 and GmZF351, which are 51.3% identical, regulate distinct genes and physically interact with each other to activate downstream genes (Lu et al., 2021). Motivated by cases like this, we selected a total of 284 TFs within oil-related regions. We used TFs found in at least two of the three studies mentioned earlier (Moharana & Venancio, 2020; Niu et al., 2020; Yao et al., 2020) (Supplementary table S5) to investigate the TF interaction patterns in the STRING database (Szklarczyk et al., 2017).

From the 284 selected TFs, 31 had interactions in STRING, of which 35.48%, 29.03% and 9.68% were up-regulated in high oil content accessions of soybean, detected in GWAS or both, respectively (Supplementary table S5). GmZF392 was found in only one of the three TF datasets explored here, but was kept in the analysis based on the strong experimental evidence supporting its role in oil biosynthesis. Therefore, Figure 3 shows 32 TFs clustered in eight groups. Groups 1,2,3, and 8 showed genes upregulated in high oil content accessions and will be discussed below.

**Figure 3.**
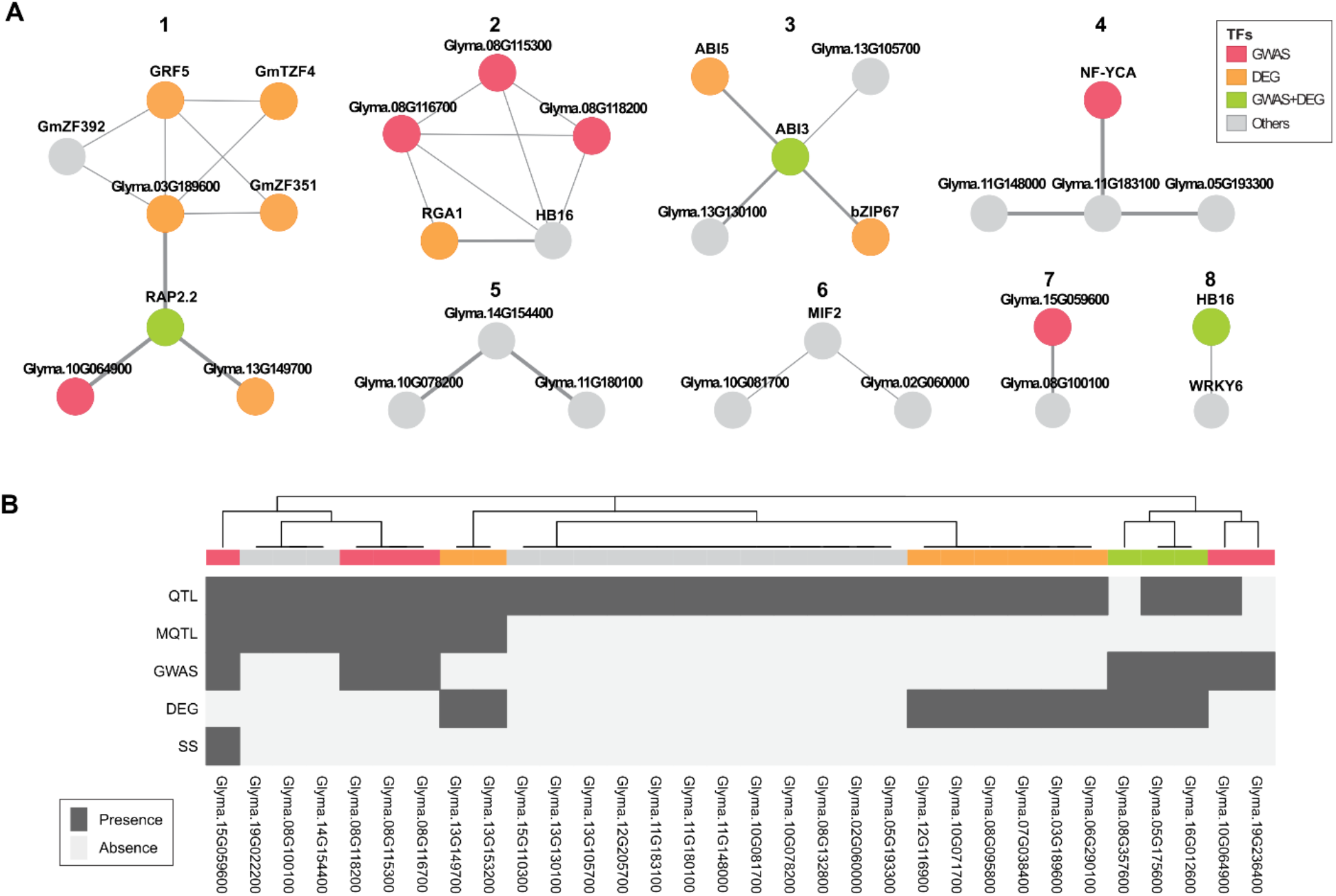
Interaction analysis candidate transcription factors (TFs). **A**. The TFs in eight interaction groups constructed using the STRING database (Szklarczyk et al., 2017). These associations refer to physical or functional relationships with a high confidence interaction score (0.7), where the edge weight is proportional to the support (Supplementary table S6). Nodes were colored as follows: red (genes present GWAS regions); orange (differentially expressed gene – DEG – upregulated when comparing high and low oil content soybean accessions); green (genes reported in both GWAS and DEG); gray (genes previously reported in quantitative trait loci [QTL], meta-QTL [MQTL] or selective sweep regions [SS]). **B**. Presence/absence profiles of the 32 candidate TFs in genomic regions for oil-related traits. The bar above the heatmap was colored as described in panel A.

Group 1 contains eight TFs from the families: GRF (Glyma.07G038400 [GRF5]), ERF (Glyma.16G012600 [RAP2.2]), Trihelix (Glyma.03G189600; Glyma.10G064900; Glyma.13G149700), and C3H (Glyma.06G290100 [GmZF351]; Glyma.12G116900 [GmTZF4]; Glyma.12G205700 [GmZF392]). Interestingly, GmZF351 and GmZF392 showed the same interactions in the network (Figure 3A). From the eight genes in Group 1, six were upregulated in high oil content accessions (Figure 3B). GRF5, Glyma.03G189600 and GmTZF4 were previously predicted to be candidates in regulation of seed lipid biosynthesis (Niu et al., 2020; Niu et al., 2020; Zhang et al., 2016). Group 2 shows associations among five TFs from the families GRAS (Glyma.08G095800 [RGA1]), HD-ZIP (Glyma.08G132800 [HB16]), bZIP (Glyma.08G115300), G2-like (Glyma.08G116700), and WRKY (Glyma.08G118200). The relationship between these TFs in the regulation of oil biosynthesis unclear, although mutants of RGA1 – a negative regulator in the gibberellin signaling pathway – have alterations in seed fatty acid metabolism in *Arabidopsis* (Chen et al., 2012). Group 3 shows associations among five TFs from the family B3 (Glyma.08G357600 [ABI3]), bZIP (Glyma.10G071700 [ABI5]; Glyma.13G153200 [bZIP67]), HSF (Glyma.13G105700), and bHLH (Glyma.13G130100). Among them, ABI3, ABI5, and bZIP67 were found to be involved in oil biosynthesis (Mendes et al., 2013; Zhang et al., 2016; Zhang et al., 2017). Group 8 shows the association between Glyma.05G175600 (HD-ZIP family, HB16) and Glyma.15G110300 (WRKY family, WRKY6). WRKY6 was downregulated in high oil content accessions (Niu et al., 2020). We believe that interaction maps like the one presented here, integrating various sources of evidence, can help us understand the regulatory dynamics involved in oil biosynthesis and in the discovery of new potential regulators.

### Identification of candidate genes for oil-related traits

Since the first QTL study of oil-related traits in soybean (Diers et al., 1992), a large set of genomic data have accumulated, including hundreds QTL and GWAS regions, as well as loci that underwent positive selection during domestication and/or breeding (Zhou et al., 2015), as discussed above. As demonstrated in this work, integrating such data can help mitigate the individual limitations of each technique. For example, QTL mapping is efficient in finding rare genes of large effect in artificial populations, but only alleles that segregate between the F2 and its progeny can be assessed (Mackay et al., 2009). On the other hand, GWAS work well with natural populations, but is limited in detecting rare alleles in a population (Korte & Farlow, 2013). Furthermore, identifying domestication or breeding selective sweep regions may also unveil powerful candidate genes (Han et al., 2016). We integrated the positional information from MQTL, GWAS and selective sweep regions to better understand the context of oil-related genes therein. These datasets contain 11,403, 6,229, and 2,230 genes, respectively (Supplementary table S1; Supplementary figure S2). In total, we found 157 genes co-located in MQTL, GWAS, and selective sweep regions (Supplementary table S7; Supplementary figure S4). Remarkably, 33.75% (53/157) of these genes are within hotspot regions for oil and protein (Qi et al., 2018), i.e. those reported in at least four different soybean QTL studies related with both, oil and protein content. The 157 genes are located in well defined regions of Gm08 (8.61 – 8.71 Mb), Gm14 (34.25 – 34.74 Mb and 45.0 – 45.2 Mb), Gm15 (4.47 – 5.17 Mb and 11.06 – 11.12 Mb), Gm16 (4.36 – 4.53 Mb), and Gm19 (41.94 – 42.32 Mb) (Figure 4). Importantly, these hotspot regions comprise only 1.949 genes (Qi et al., 2018).

**Figure 4.**
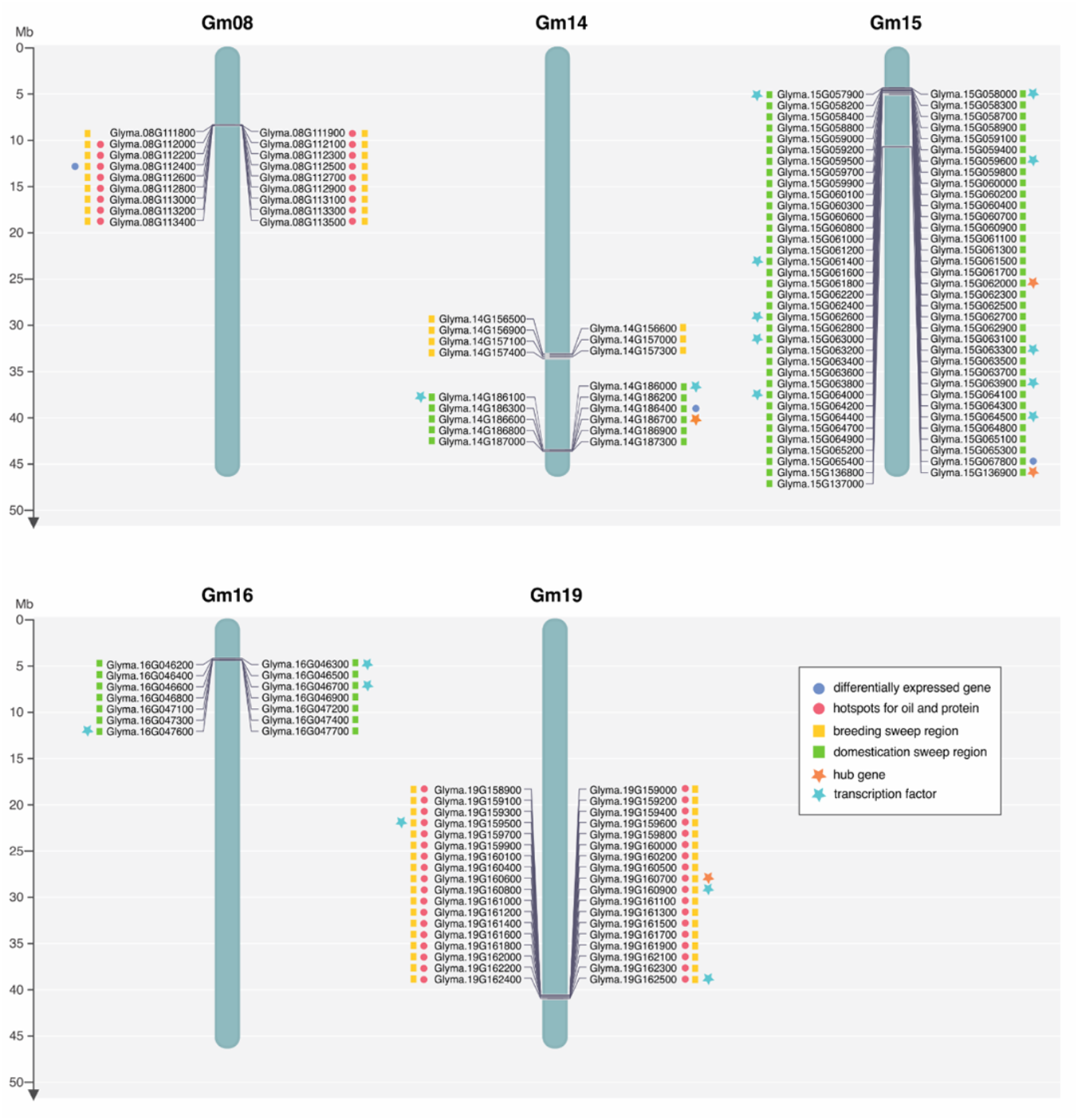
Chromosome map with 157 genes co-located in MQTL, GWAS, and selective sweep regions. Differentially expressed genes were obtained by a comparison of high and low oil content soybean accessions (Niu et al., 2020). Hotspots for oil and protein and selective sweep regions were reported by Qi et al. (2018) and Zhou et al. (2015), respectively. Hub genes are those with the highest number of interactions in expression modules positively correlated with oil-related traits (Qi et al., 2018; Yang et al., 2019). More details on the annotations of these genes are provided in Supplementary table S7.

Some remarkable trends emerge from these regions. The Gm08 and Gm19 regions harbor breeding sweep regions and are associated with oil and protein content. Similarly, all the Gm15 and Gm16 blocks corresponded only to domestication sweep regions. We also found that oil and protein hotspots are strongly associated with breeding and never with domestication, suggesting that domestication prioritized the selection of other traits. Interestingly, 15 of 18 TFs found among the 157 genes within these blocks are associated with domestication. Together, these results indicate that domestication involved the selection of regulatory programs, whereas breeding appears to involve mainly lipid and other metabolic genes (Supplementary table S8). It is also conceivable that several TFs selected during domestication regulate transcriptional programs that were important for breeding.

We further investigated the functions of the 157 genes within these regions using gene coexpression networks from two studies, in particular with regard to the hubs from oil-related modules. Qi et al. found 96 hub genes in a module (brown) positively correlated with oil-related traits (Qi et al., 2018). Of these, four (i.e. Glyma.14G186700, Glyma.15G062000, Glyma.19G160700, and Glyma.15G136900) are among the 157 genes reported above (Figure 4; Supplementary table S9; Supplementary figure S5). These genes are likely involved in defense/immunity (Glyma.14G186700; Glyma.15G062000) or regulatory processes (Glyma.19G160700; Glyma.15G136900). The only hub gene in a breeding region is Glyma.19G160700 (an ortholog of AT3G54610), which encode a GNAT histone acetyl transferase (GCN5). Wang et al. revealed that GCN5-dependent H3K9/14 acetylation of Omega-3 fatty acid desaturase (FAD3) determined the expression levels of FAD3 in *A. thaliana* seeds. Moreover, the ratio of linolenic to linoleic acid in the *gcn5* mutant was rescued to the wild-type levels through the overexpression of FAD3 (Wang et al., 2016). These results make Glyma.19G160700 a promising candidate to improve soybean oil quality.

Yang et al. found a total of 31 hubs in modules (pink, brown and blue) positively correlated with oil-related traits (Yang et al., 2019). Interestingly, five (16.13%) of these hubs (Glyma.03G204400, Glyma.05G234000, Glyma.06G195000, Glyma.07G196200, and Glyma.19G228800) are reported in at least two different studies considered here (Supplementary table S9; Supplementary figure S5). Among them, Glyma.05G234000 (an ortholog of AT5G16110) encodes a hypotetical protein. According to data from TAIR, although this gene is expressed in several tissues and its protein product locates to the chloroplast, there is no evidence about its molecular functions or conserved domains (Rhee et al., 2003). Glyma.19G228800 (an ortholog of AT4G02080) encodes an ADP-ribosylation factor-relate (ARF1) that can be involved in transport from the endoplasmic reticulum to the Golgi apparatus (Matheson et al., 2007). Glyma.03G204400 (an ortholog of AT5G22000) present a RING-H2 conserved domain. Previous studies have shown that RING-finger proteins are involved in plant growth and development, stress resistance, hormone signaling responses and controlling characteristics of both vegetative and seed yield (Sun et al., 2019; Zombori et al., 2020). Glyma.06G195000 (an ortholog of AT2G03090) present two expansin domains, a pollen allergen domain, and a rare lipoprotein A (RlpA)-like domain. Finally, Glyma.07G196200 (an ortholog of AT4G21610) encode a LSD1 zinc finger, involved in rice growth and disease resistance (Xu & He, 2007). The roles of these genes in oil biosynthesis are yet to be characterized.

The negative correlation between oil and protein contents in soybean has been widely reported over decades (Bandillo et al., 2015; Chaudhary et al., 2015; Johnson & Bernard, 1962; Patil et al., 2017). Hence, identifying regions that simultaneously contribute to these traits have been a topic of great interest. In hotspot regions, we found Glyma.08G112300 (an ortholog of AT1G55260) that encodes a multifunctional 2S albumin superfamily protein involved in defense and storage (Lin et al., 2004). Glyma.08G112300 has been reported as a strong determinant of high levels of water-soluble proteins, a critical factor both in food quality and in the production of isolated soybean proteins (Zhang et al., 2017). However, this gene is highly expressed across several tissues (Supplementary figure S6). Curiously, only 2kb away from Glyma.08G112300, we found Glyma.08G112400 (Figure 4; Supplementary figure S7; Supplementary table S7), which encodes a protein with a domain of unknown function DUF538 that has been recently proposed as a candidate gene to determine oleic acid content (Niu et al., 2020). The genomic closeness of these two genes indicate that they should be investigated in more detail with regard to their roles in determining protein and oil content.

## CONCLUSIONS

Here, we used integrative approaches to explore genes in stable genomic regions for oil-related traits in soybean. We explored publicly available datasets, mainly from studies of QTL, MQTL, GWAS, and selective sweep regions. This core dataset was complemented with gene expression data, gene expression networks and TF annotations to help elucidating the genetic basis of oil-related traits. The integrative analyses reported here provide a framework to identify and prioritize candidate genes. Finally, the gene set reported here might be an important repository for experimental validation and soybean improvement programs.

## MATERIALS AND METHODS

### Oil-related QTL and GWAS data

Coordinates of QTL and several genetic markers were retrieved from SoyBase (update from January 2018; soybase.org). QTL were extracted with in-house Perl and bash scripts. The flanking regions of the closest genetic markers were used to define the ends of each QTL. Next, we used a bash script to extract a total of 4,352 names of QTL objects, which had their chromosomal coordinates determined. From these, we found 478 oil-related QTL (Supplementary table S10) and retrieved the genes therein using the file Gmax_275_Wm82.a2.v1.gene.gff3, from Phytozome V12.1, as reference. The files containing the coordinates of 478 oil-related QTL and genes (gff3) were integrated in Browser Extensible Data (BED) format with Bedtools V2.27.1 (Quinlan, 2014), followed by redundancy removal (Supplementary figure S1; Table S1). The GWAS data were obtained from 15 studies (Table 1). The coordinates of the significant LD regions (Supplementary table S2), corresponding to the SNPs reported in the original studies, were collected in individual files. Six of these studies used an older version of the soybean reference genome (Glyma.Wm82.a1.v1) and had their data converted using the same reference genome file mentioned above. These files were combined and processed to remove redundancy, resulting in a GWAS list with 6,804 genes (Supplementary figure S1; Supplementary table S1).

### Lipid metabolism pathways

Information about genes in metabolic pathways were obtained from soybase.org and from Yao et al. 2020 (Supplementary table S8 (column QTL-LM); Supplementary table S11).

### MQTL, hub genes, DEGs, selective sweep and transcription factors

We supplemented our data using MQTL, hub genes from modules positively correlated with oil, DEGs from soybean accessions of low to high oil content, positively selected (selective sweep) regions during seed domestication/breeding and TF classifications. The chromosomal map with genes co-located in selective sweeps, MQTL, and GWAS regions was constructed using MapGene2Chrom (Jiangtao et al., 2015). A workflow is available at Supplementary figure S2.

The coordinates of MQTL were obtained from Qi et al. (Qi et al., 2018). We retrieved 63, 26 and 11 MQTL for oil content, fatty acid and hotspots for oil and protein, respectively (Supplementary tables S12 and S13). MQTL coordinates and genes were retrieved with the same strategy used for QTL. We also used hubs from a coexpression modules reported by two previous studies (Supplementary table S8) (Qi et al., 2018; Yang et al., 2019).

We used DEGs from two publications. Yang et al. reported DEGs between 20 and 10 days after flowering – a critical seed oil accumulation stage in the soybean variety ‘nannong1138-2’ (NN1138-2), which shows ∼19% seed oil content (Yang et al., 2019). The second dataset, from Niu et al. encompasses six soybean accessions with ∼11% to 22% seed oil content (Niu et al., 2020). Selective sweep regions (Zhou et al., 2015) were retrieved and had their gene names/coordinates updated as described above for the GWAS (Supplementary table S8).

TFs were recovered from three sources. TFs reported in stable QTL regions (Yao et al., 2020); TFs differentially expressed in soybean accessions with divergent oil content and compositions (Niu et al., 2020); and from a systematic classification of legume TF repertoires (Moharana & Venancio, 2020). These data are available in Supplementary table S8. TF interaction analysis was conducted using STRING version 11 (Szklarczyk et al., 2017), using a 0.7 (high) confidence threshold. To the input we used the file Gmax_275_Wm82.a2.v1.protein.fa. from Phytozome V12.1, as reference.

### Global gene expression

Analyses of global expression genes were conducted from Soybean Expression Atlas database (Machado et al., 2020), using Kallisto as a method for estimating gene expression.

## ACKNOWLEDGEMENTS

This work was supported by Fundação Carlos Chagas Filho de Amparo à Pesquisa do Estado do Rio de Janeiro (FAPERJ; grants E-26/203.309/2016 and E-26/203.014/2018), Coordenação de Aperfeiçoamento de Pessoal de Nível Superior - Brasil (CAPES; Finance Code 001), and Conselho Nacional de Desenvolvimento Científico e Tecnológico. The funding agencies had no role in the design of the study and collection, analysis, and interpretation of data and in writing.

## REFERENCES

Abdelghany, A. M., Zhang, S., Azam, M., Shaibu, A. S., Feng, Y., Qi, J., Li, Y., Tian, Y., Hong, H., Li, B., & Sun, J. (2020). Natural variation in fatty acid composition of diverse world soybean germplasms grown in China. Agronomy, 10(1), 1–18. https://doi.org/10.3390/agronomy10010024

Akond, M., Liu, S., Boney, M., Kantartzi, S. K., Meksem, K., Bellaloui, N., Lightfoot, D. A., & Kassem, M. A. (2014). Identification of Quantitative Trait Loci (QTL) Underlying Protein, Oil, and Five Major Fatty Acids’ Contents in Soybean. American Journal of Plant Sciences, 05(01), 158–167. https://doi.org/10.4236/ajps.2014.51021

Almeida-Silva, F., Moharana, K. C., Machado, F. B., & Venancio, T. M. (2020). Exploring the complexity of soybean (Glycine max) transcriptional regulation using global gene co-expression networks. Planta, (in press). https://doi.org/10.1007/s00425-020-03499-8

Almeida-Silva, Fabricio, Moharana, K. C., & Venancio, T. M. (2021). The state of the art in soybean transcriptomics resources and gene coexpression networks. In Silico Plants, 3(1), 1–7. https://doi.org/10.1093/insilicoplants/diab005

Bachlava, E., Dewey, R. E., Burton, J. W., & Cardinal, A. J. (2009). Mapping and comparison of quantitative trait loci for oleic acid seed content in two segregating soybean populations. Crop Science, 49(2), 433–442. https://doi.org/10.2135/cropsci2008.06.0324

Bandillo, N., Jarquin, D., Song, Q., Nelson, R., Cregan, P., Specht, J., & Lorenz, A. (2015). A Population Structure and Genome-Wide Association Analysis on the USDA Soybean Germplasm Collection. The Plant Genome, 8(3), 0. https://doi.org/10.3835/plantgenome2015.04.0024

Bates, P. D., Stymne, S., & Ohlrogge, J. (2013). Biochemical pathways in seed oil synthesis. Current Opinion in Plant Biology, 16(3), 358–364. https://doi.org/10.1016/j.pbi.2013.02.015

Bellieny-Rabelo, D., De Oliveira, E. A. G., Da Silva Ribeiro, E., Pessoa Costa, E., Oliveira, A. E. A., & Venancio, T. M. (2016). Transcriptome analysis uncovers key regulatory and metabolic aspects of soybean embryonic axes during germination. Scientific Reports, 6 (October), 1–12. https://doi.org/10.1038/srep36009

Borevitz, J. O., & Nordborg, M. (2003). Update on genomics and natural variation in Arabidopsis: The impact of genomics on the study of natural variation in Arabidopsis. Plant Physiology, 132 (June), 718–725. https://doi.org/10.1104/pp.103.023549.718

Bourgis, F., Kilaru, A., Cao, X., Ngando-Ebongue, G.-F., Drira, N., Ohlrogge, J. B., & Arondel, V. (2011). Comparative transcriptome and metabolite analysis of oil palm and date palm mesocarp that differ dramatically in carbon partitioning. Proceedings of the National Academy of Sciences, 108(30), 12527–12532. https://doi.org/10.1073/pnas.1106502108

Cao, Y., Li, S., Wang, Z., Chang, F., Kong, J., Gai, J., & Zhao, T. (2017). Identification of Major Quantitative Trait Loci for Seed Oil Content in Soybeans by Combining Linkage and Genome-Wide Association Mapping. Frontiers in Plant Science, 8(July), 1–10. https://doi.org/10.3389/fpls.2017.01222

Chaudhary, J., Patil, G. B., Sonah, H., Deshmukh, R. K., Vuong, T. D., Valliyodan, B., & Nguyen, H. T. (2015). Expanding Omics Resources for Improvement of Soybean Seed Composition Traits. Frontiers in Plant Science, 6(November), 1–16. https://doi.org/10.3389/fpls.2015.01021

Chen, C., Meng, Y., Shopan, J., Whelan, J., Hu, Z., Yang, J., & Zhang, M. (2020). Identification and characterization of Arabidopsis thaliana mitochondrial F1F0-ATPase inhibitor factor 1. Journal of Plant Physiology, 254(March), 153264. https://doi.org/10.1016/j.jplph.2020.153264

Chen, H., Patterson, N., & Reich, D. (2010). Population differentiation as a test for selective sweeps. Genome Research, 20(3), 393–402. https://doi.org/10.1101/gr.100545.109

Chen, M., Du, X., Zhu, Y., Wang, Z., Hua, S., Li, Z., Guo, W., Zhang, G., Peng, J., & Jiang, L. (2012). Seed Fatty Acid Reducer acts downstream of gibberellin signalling pathway to lower seed fatty acid storage in Arabidopsis. Plant, Cell and Environment, 35(12), 2155–2169. https://doi.org/10.1111/j.1365-3040.2012.02546.x

Clemente, T. E., & Cahoon, E. B. (2009). Soybean Oil: Genetic Approaches for Modification of Functionality and Total Content. Plant Physiology, 151(3), 1030–1040. https://doi.org/10.1104/pp.109.146282

Collard, B. C. Y., & Mackill, D. J. (2008). Marker-assisted selection: An approach for precision plant breeding in the twenty-first century. Philosophical Transactions of the Royal Society B: Biological Sciences, 363(1491), 557–572. https://doi.org/10.1098/rstb.2007.2170

Daware, A. V., Srivastava, R., Singh, A. K., Parida, S. K., & Tyagi, A. K. (2017). Regional Association Analysis of MetaQTLs Delineates Candidate Grain Size Genes in Rice. Frontiers in Plant Science, 8(May). https://doi.org/10.3389/fpls.2017.00807

Diers, B. W., Keim, P., Fehr, W. R., & Shoemaker, R. C. (1992). RFLP analysis of soybean seed protein and oil content. Theoretical and Applied Genetics, 83(5), 608–612. https://doi.org/10.1007/BF00226905

Eskandari, M., Cober, E. R., & Rajcan, I. (2013). Genetic control of soybean seed oil: II. QTL and genes that increase oil concentration without decreasing protein or with increased seed yield. Theoretical and Applied Genetics, 126(6), 1677–1687. https://doi.org/10.1007/s00122-013-2083-z

Fang, C., Ma, Y., Wu, S., Liu, Z., Wang, Z., Yang, R., Hu, G., Zhou, Z., Yu, H., Zhang, M., Pan, Y., Zhou, G., Ren, H., Du, W., Yan, H., Wang, Y., Han, D., Shen, Y., Liu, S., … Tian, Z. (2017). Genome-wide association studies dissect the genetic networks underlying agronomical traits in soybean. Genome Biology, 18(1), 161. https://doi.org/10.1186/s13059-017-1289-9

Goffinet, B., & Gerber, S. (2000). Quantitative trait loci: A meta-analysis. Genetics, 155(1), 463–473.

Gong, Q. chun, Yu, H. xiao, Mao, X. rui, Qi, H. dong, Shi, Y., Xiang, W., Chen, Q. shan, & Qi, Z. ming. (2018). Meta-analysis of soybean amino acid QTLs and candidate gene mining. Journal of Integrative Agriculture, 17(5), 1074–1084. https://doi.org/10.1016/S2095-3119(17)61783-0

Guillaumot, D., Guillon, S., Déplanque, T., Vanhee, C., Gumy, C., Masquelier, D., Morsomme, P., & Batoko, H. (2009). The Arabidopsis TSPO-related protein is a stress and abscisic acid-regulated, endoplasmic reticulum-Golgi-localized membrane protein. Plant Journal, 60(2), 242–256. https://doi.org/10.1111/j.1365-313X.2009.03950.x

Guillaumot, D., Guillon, S., Morsomme, P., & Batoko, H. (2009). ABA, porphyrins and plant TSPO-related protein. Plant Signaling & Behavior, 4(11), 1087–1090. https://doi.org/10.4161/psb.4.11.9796

Gupta, M., Bhaskar, P. B., Sriram, S., & Wang, P.-H. (2017). Integration of omics approaches to understand oil/protein content during seed development in oilseed crops. Plant Cell Reports, 36(5), 637–652. https://doi.org/10.1007/s00299-016-2064-1

Han, Y., Zhao, X., Liu, D., Li, Y., Lightfoot, D. A., Yang, Z., Zhao, L., Zhou, G., Wang, Z., Huang, L., Zhang, Z., Qiu, L., Zheng, H., & Li, W. (2016). Domestication footprints anchor genomic regions of agronomic importance in soybeans. New Phytologist, 209(2), 871–884. https://doi.org/10.1111/nph.13626

Haun, W., Coffman, A., Clasen, B. M., Demorest, Z. L., Lowy, A., Ray, E., Retterath, A., Stoddard, T., Juillerat, A., Cedrone, F., Mathis, L., Voytas, D. F., & Zhang, F. (2014). Improved soybean oil quality by targeted mutagenesis of the fatty acid desaturase 2 gene family. Plant Biotechnology Journal, 12(7), 934–940. https://doi.org/10.1111/pbi.12201

Hwang, E. Y., Song, Q., Jia, G., Specht, J. E., Hyten, D. L., Costa, J., & Cregan, P. B. (2014). A genome-wide association study of seed protein and oil content in soybean. BMC Genomics, 15(1), 1–12. https://doi.org/10.1186/1471-2164-15-1

Hyten, D. L., Song, Q., Zhu, Y., Choi, I. Y., Nelson, R. L., Costa, J. M., Specht, J. E., Shoemaker, R. C., & Cregan, P. B. (2006). Impact of genetic bottlenecks on soybean genome diversity. Proceedings of the National Academy of Sciences of the United States of America, 103(45), 16666–16671. https://doi.org/10.1073/pnas.0604379103

Jiangtao, C., Yingzhen, K., Qian, W., Yuhe, S., Daping, G., Jing, L., & Guanshan, L. (2015). MapGene2Chrom, a tool to draw gene physical map based on Perl and SVG languages. Yi Chuan = Hereditas, 37(1), 91–97. https://doi.org/10.16288/j.yczz.2015.01.013

Johnson, H. W., & Bernard, R. L. (1962). Soybean Genetics And Breeding. In Journal of Chemical Information and Modeling (Vol. 53, Issue 9, pp. 149–221). https://doi.org/10.1016/S0065-2113(08)60438-1

Jurkiewicz, P., Melser, S., Maucourt, M., Ayeb, H., Veljanovski, V., Maneta-Peyret, L., Hooks, M., Rolin, D., Moreau, P., & Batoko, H. (2018). The multistress-induced Translocator protein (TSPO) differentially modulates storage lipids metabolism in seeds and seedlings. The Plant Journal, 96(2), 274–286. https://doi.org/10.1111/tpj.14028

Kanai, M., Yamada, T., Hayashi, M., Mano, S., & Nishimura, M. (2019). Soybean (Glycine max L.) triacylglycerol lipase GmSDP1 regulates the quality and quantity of seed oil. Scientific Reports, 9(1), 1–10. https://doi.org/10.1038/s41598-019-45331-8

Karikari, B., Li, S., Bhat, J., Cao, Y., Kong, J., Yang, J., Gai, J., & Zhao, T. (2019). Genome-Wide Detection of Major and Epistatic Effect QTLs for Seed Protein and Oil Content in Soybean Under Multiple Environments Using High-Density Bin Map. International Journal of Molecular Sciences, 20(4), 979. https://doi.org/10.3390/ijms20040979

Kong, Q., Singh, S. K., Mantyla, J. J., Pattanaik, S., Guo, L., Yuan, L., Benning, C., & Ma, W. (2020). Teosinte branched1/cycloidea/ proliferating cell factor4 interacts with wrinkled1 to mediate seed oil biosynthesis. Plant Physiology, 184(2), 658–665. https://doi.org/10.1104/pp.20.00547

Koo, A. J. K., Ohlrogge, J. B., & Pollard, M. (2004). On the Export of Fatty Acids from the Chloroplast. Journal of Biological Chemistry, 279(16), 16101–16110. https://doi.org/10.1074/jbc.M311305200

Korte, A., & Farlow, A. (2013). The advantages and limitations of trait analysis awith GWAS: a review. Plant Methods, 9(29), 1–9.

Leamy, L. J., Zhang, H., Li, C., Chen, C. Y., & Song, B.-H. (2017). A genome-wide association study of seed composition traits in wild soybean (Glycine soja). BMC Genomics, 18(1), 18. https://doi.org/10.1186/s12864-016-3397-4

Li, D., Zhao, X., Han, Y., Li, W., & Xie, F. (2019). Genome-wide association mapping for seed protein and oil contents using a large panel of soybean accessions. Genomics, 111(1), 90–95. https://doi.org/10.1016/j.ygeno.2018.01.004

Li, N., Xu, C., Li-Beisson, Y., & Philippar, K. (2016). Fatty Acid and Lipid Transport in Plant Cells. Trends in Plant Science, 21(2), 145–158. https://doi.org/10.1016/j.tplants.2015.10.011

Li, Q.-T., Lu, X., Song, Q.-X., Chen, H.-W., Wei, W., Tao, J.-J., Bian, X.-H., Shen, M., Ma, B., Zhang, W.-K., Bi, Y.-D., Li, W., Lai, Y.-C., Lam, S.-M., Shui, G.-H., Chen, S.-Y., & Zhang, J.-S. (2017). Selection for a Zinc-Finger Protein Contributes to Seed Oil Increase during Soybean Domestication. Plant Physiology, 173(4), 2208–2224. https://doi.org/10.1104/pp.16.01610

Li, Xu, Yang, & Zhao. (2019). Dissecting the Genetic Architecture of Seed Protein and Oil Content in Soybean from the Yangtze and Huaihe River Valleys Using Multi-Locus Genome-Wide Association Studies. International Journal of Molecular Sciences, 20(12), 3041. https://doi.org/10.3390/ijms20123041

Li, Y., Reif, J. C., Hong, H., Li, H., Liu, Z., Ma, Y., Li, J., Tian, Y., Li, Y., Li, W., & Qiu, L. (2018). Genome-wide association mapping of QTL underlying seed oil and protein contents of a diverse panel of soybean accessions. Plant Science, 266, 95–101. https://doi.org/10.1016/j.plantsci.2017.04.013

Li, Y., Reif, J. C., Ma, Y., Hong, H., Liu, Z., Chang, R., & Qiu, L. (2015). Targeted association mapping demonstrating the complex molecular genetics of fatty acid formation in soybean. BMC Genomics, 16(1), 841. https://doi.org/10.1186/s12864-015-2049-4

Lin, J., Fido, R., Shewry, P., Archer, D. B., & Alcocer, M. J. C. (2004). The expression and processing of two recombinant 2S albumins from soybean (Glycine max) in the yeast Pichia pastoris. Biochimica et Biophysica Acta (BBA) - Proteins and Proteomics, 1698(2), 203–212. https://doi.org/10.1016/j.bbapap.2003.12.001

Lin, L.-J., Tai, S. S. K., Peng, C.-C., & Tzen, J. T. C. (2002). Steroleosin, a sterol-binding dehydrogenase in seed oil bodies. Plant Physiology, 128(4), 1200–1211. https://doi.org/10.1104/pp.010928

Liu, Li P., Zhang, Y. W., Zuo, J. F., Li, G., Han, X., Dunwell, J. M., & Zhang, Y. M. (2020). Three-dimensional genetic networks among seed oil-related traits, metabolites and genes reveal the genetic foundations of oil synthesis in soybean. Plant Journal, 103(3), 1103–1124. https://doi.org/10.1111/tpj.14788

Liu, Y., Du, H., Li, P., Shen, Y., Peng, H., Liu, S., Zhou, G.-A., Zhang, H., Liu, Z., Shi, M., Huang, X., Li, Y., Zhang, M., Wang, Z., Zhu, B., Han, B., Liang, C., & Tian, Z. (2020). Pan-Genome of Wild and Cultivated Soybeans. Cell, 0(0), 1–15. https://doi.org/10.1016/j.cell.2020.05.023

Liu, Y. F., Li, Q. T., Lu, X., Song, Q. X., Lam, S. M., Zhang, W. K., Ma, B., Lin, Q., Man, W. Q., Du, W. G., Shui, G. H., Chen, S. Y., & Zhang, J. S. (2014). Soybean GmMYB73 promotes lipid accumulation in transgenic plants. BMC Plant Biology, 14(1), 1–16. https://doi.org/10.1186/1471-2229-14-73

Lu, L., Wei, W., Li, Q., Bian, X., Lu, X., Hu, Y., Cheng, T., Wang, Z., Jin, M., Tao, J., Yin, C., He, S., Man, W., Li, W., Lai, Y., Zhang, W., Chen, S., & Zhang, J. (2021). A transcriptional regulatory module controls lipid accumulation in soybean. In New Phytologist. https://doi.org/10.1111/nph.17401

Lu, X., Li, Q. T., Xiong, Q., Li, W., Bi, Y. D., Lai, Y. C., Liu, X. L., Man, W. Q., Zhang, W. K., Ma, B., Chen, S. Y., & Zhang, J. S. (2016). The transcriptomic signature of developing soybean seeds reveals the genetic basis of seed trait adaptation during domestication. The Plant Journal : For Cell and Molecular Biology, 86(6), 530–544. https://doi.org/10.1111/tpj.13181

Machado, F. B., Moharana, K. C., Almeida-Silva, F., Gazara, R. K., Pedrosa-Silva, F., Coelho, F. S., Grativol, C., & Venancio, T. M. (2020). Systematic analysis of 1,298 RNA-Seq samples and construction of a comprehensive soybean (Glycine max) expression atlas. The Plant Journal, tpj.14850. https://doi.org/10.1111/tpj.14850

Mackay, T. F. C., Stone, E. A., & Ayroles, J. F. (2009). The genetics of quantitative traits: challenges and prospects. Nature Reviews Genetics, 10(8), 565–577. https://doi.org/10.1038/nrg2612

Manan, S., Chen, B., She, G., Wan, X., & Zhao, J. (2017). Transport and transcriptional regulation of oil production in plants. Critical Reviews in Biotechnology, 37(5), 641–655. https://doi.org/10.1080/07388551.2016.1212185

Marchive, C., Nikovics, K., To, A., Lepiniec, L., & Baud, S. (2014). Transcriptional regulation of fatty acid production in higher plants: Molecular bases and biotechnological outcomes. European Journal of Lipid Science and Technology, 116(10), 1332–1343. https://doi.org/10.1002/ejlt.201400027

Matheson, L. A., Hanton, S. L., Rossi, M., Latijnhouwers, M., Stefano, G., Renna, L., & Brandizzi, F. (2007). Multiple roles of ADP-ribosylation factor 1 in plant cells include spatially regulated recruitment of coatomer and elements of the Golgi matrix. Plant Physiology, 143(4), 1615–1627. https://doi.org/10.1104/pp.106.094953

Mendes, A., Kelly, A. A., van Erp, H., Shaw, E., Powers, S. J., Kurup, S., & Eastmond, P. J. (2013). bZIP67 regulates the omega-3 fatty acid content of arabidopsis seed oil by activating fatty acid DESATURASE3. Plant Cell, 25(8), 3104–3116. https://doi.org/10.1105/tpc.113.116343

Moharana, K. C., & Venancio, T. M. (2020). Polyploidization events shaped the transcription factor repertoires in legumes (Fabaceae). The Plant Journal, 103(2), 726–741. https://doi.org/10.1111/tpj.14765

Niu, Yuan, Zhang, G., Wan, F., & Zhang, Y.-M. (2020). Integration of RNA-Seq profiling with genome-wide association study predicts candidate genes for oil accumulation in soybean. Crop and Pasture Science, 71(12), 996. https://doi.org/10.1071/CP20358

Niu, Yue, Wu, L., Li, Y., Huang, H., Qian, M., Sun, W., Zhu, H., Xu, Y., Fan, Y., Mahmood, U., Xu, B., Zhang, K., Qu, C., Li, J., & Lu, K. (2020). Deciphering the transcriptional regulatory networks that control size, color, and oil content in Brassica rapa seeds. Biotechnology for Biofuels, 13(1), 1–20. https://doi.org/10.1186/s13068-020-01728-6

Patil, G., Mian, R., Vuong, T., Pantalone, V., Song, Q., Chen, P., Shannon, G. J., Carter, T. C., & Nguyen, H. T. (2017). Molecular mapping and genomics of soybean seed protein: a review and perspective for the future. Theoretical and Applied Genetics, 130(10), 1975–1991. https://doi.org/10.1007/s00122-017-2955-8

Patil, G., Vuong, T. D., Kale, S., Valliyodan, B., Deshmukh, R., Zhu, C., Wu, X., Bai, Y., Yungbluth, D., Lu, F., Kumpatla, S., Shannon, J. G., Varshney, R. K., & Nguyen, H. T. (2018). Dissecting genomic hotspots underlying seed protein, oil, and sucrose content in an interspecific mapping population of soybean using high-density linkage mapping. Plant Biotechnology Journal, 16(11), 1939–1953. https://doi.org/10.1111/pbi.12929

Pham, A. T., Shannon, J. G., & Bilyeu, K. D. (2012). Combinations of mutant FAD2 and FAD3 genes to produce high oleic acid and low linolenic acid soybean oil. Theoretical and Applied Genetics, 125(3), 503–515. https://doi.org/10.1007/s00122-012-1849-z

Priolli, R. H. G., Campos, J. B., Stabellini, N. S., Pinheiro, J. B., & Vello, N. A. (2015). Association mapping of oil content and fatty acid components in soybean. Euphytica, 203(1), 83–96. https://doi.org/10.1007/s10681-014-1264-4

Qi, Zhao-ming, Wu, Q., Han, X., Sun, Y., Du, X., Liu, C., Jiang, H., Hu, G., & Chen, Q. (2011). Soybean oil content QTL mapping and integrating with meta-analysis method for mining genes. Euphytica, 179(3), 499–514. https://doi.org/10.1007/s10681-011-0386-1

Qi, Zhaoming, Zhang, Z., Wang, Z., Yu, J., Qin, H., Mao, X., Jiang, H., Xin, D., Yin, Z., Zhu, R., Liu, C., Yu, W., Hu, Z., Wu, X., Liu, J., & Chen, Q. (2018). Meta-analysis and transcriptome profiling reveal hub genes for soybean seed storage composition during seed development. Plant, Cell & Environment, 41(9), 2109–2127. https://doi.org/10.1111/pce.13175

Qin, H., Liu, Z., Wang, Y., Xu, M., Mao, X., Qi, H., Yin, Z., Li, Y., Jiang, H., Hu, Z., Wu, X., Zhu, R., Liu, C., Chen, Q., Xin, D., & Qi, Z. (2018). Meta-analysis and overview analysis of quantitative trait locis associated with fatty acid content in soybean for candidate gene mining. Plant Breeding, 137(2), 181–193. https://doi.org/10.1111/pbr.12562

Quinlan, A. R. (2014). BEDTools: The Swiss-Army Tool for Genome Feature Analysis. Current Protocols in Bioinformatics, 47(1), 11.12.1-11.12.34. https://doi.org/10.1002/0471250953.bi1112s47

Rhee, S. Y., Beavis, W., Berardini, T. Z., Chen, G., Dixon, D., Doyle, A., Garcia-Hernandez, M., Huala, E., Lander, G., Montoya, M., Miller, N., Mueller, L. A., Mundodi, S., Reiser, L., Tacklind, J., Weems, D. C., Wu, Y., Xu, I., Yoo, D., … Zhang, P. (2003). The Arabidopsis Information Resource (TAIR): A model organism database providing a centralized, curated gateway to Arabidopsis biology, research materials and community. Nucleic Acids Research, 31(1), 224–228. https://doi.org/10.1093/nar/gkg076

Ronne, M., Labbé, C., Lebreton, A., Sonah, H., Deshmukh, R., Jean, M., Belzile, F., O’Donoughue, L., & Bélanger, R. (2019). Integrated QTL mapping, gene expression and nucleotide variation analyses to investigate complex quantitative traits: a case study with the soybean– Phytophthora sojae interaction. Plant Biotechnology Journal, pbi.13301. https://doi.org/10.1111/pbi.13301

Sacks, F. M., Lichtenstein, A. H., Wu, J. H. Y., Appel, L. J., Creager, M. A., Kris-Etherton, P. M., Miller, M., Rimm, E. B., Rudel, L. L., Robinson, J. G., Stone, N. J., & Van Horn, L. V. (2017). Dietary fats and cardiovascular disease: A presidential advisory from the American Heart Association. Circulation, 136(3), e1–e23. https://doi.org/10.1161/CIR.0000000000000510

Salie, M. J., & Thelen, J. J. (2016a). Regulation and structure of the heteromeric acetyl-CoA carboxylase. Biochimica et Biophysica Acta (BBA) - Molecular and Cell Biology of Lipids, 1861(9), 1207–1213. https://doi.org/10.1016/j.bbalip.2016.04.004

Salie, M. J., & Thelen, J. J. (2016b). Regulation and structure of the heteromeric acetyl-CoA carboxylase. Biochimica et Biophysica Acta - Molecular and Cell Biology of Lipids, 1861(9), 1207–1213. https://doi.org/10.1016/j.bbalip.2016.04.004

Sandhu, D., Alt, J. L., Scherder, C. W., Fehr, W. R., & Bhattacharyya, M. K. (2007). Enhanced oleic acid content in the soybean mutant M23 is associated with the deletion in the Fad2-1a gene encoding a fatty acid desaturase. JAOCS, Journal of the American Oil Chemists’ Society, 84(3), 229–235. https://doi.org/10.1007/s11746-007-1037-5

Schmidt, M. A., & Herman, E. M. (2008). Suppression of Soybean Oleosin Produces Micro-Oil Bodies that Aggregate into Oil Body/ER Complexes. Molecular Plant, 1(6), 910–924. https://doi.org/10.1093/mp/ssn049

Schmutz, J., Cannon, S. B., Schlueter, J., Ma, J., Mitros, T., Nelson, W., Hyten, D. L., Song, Q., Thelen, J. J., Cheng, J., Xu, D., Hellsten, U., May, G. D., Yu, Y., Sakurai, T., Umezawa, T., Bhattacharyya, M. K., Sandhu, D., Valliyodan, B., … Jackson, S. A. (2010). Genome sequence of the palaeopolyploid soybean. Nature, 463(7278), 178–183. https://doi.org/10.1038/nature08670

Sonah, H., O’Donoughue, L., Cober, E., Rajcan, I., & Belzile, F. (2015). Identification of loci governing eight agronomic traits using a GBS-GWAS approach and validation by QTL mapping in soya bean. Plant Biotechnology Journal, 13(2), 211–221. https://doi.org/10.1111/pbi.12249

Song, Q., Hyten, D. L., Jia, G., Quigley, C. V., Fickus, E. W., Nelson, R. L., & Cregan, P. B. (2013). Development and Evaluation of SoySNP50K, a High-Density Genotyping Array for Soybean. PLoS ONE, 8(1), 1–12. https://doi.org/10.1371/journal.pone.0054985 https://doi.org/10.1371/journal.pone.0054985

Song, Q., Jenkins, J., Jia, G., Hyten, D. L., Pantalone, V., Jackson, S. A., Schmutz, J., & Cregan, P. B. (2016). Construction of high resolution genetic linkage maps to improve the soybean genome sequence assembly Glyma1.01. BMC Genomics, 17(1). https://doi.org/10.1186/s12864-015-2344-0

Stephan, W. (2019). Selective Sweeps. Genetics, 211(1), 5–13. https://doi.org/10.1534/genetics.118.301319

Sun, J., Sun, Y., Ahmed, R. I., Ren, A., & Xie, M. (2019). Research progress on plant RING-finger proteins. Genes, 10(12). https://doi.org/10.3390/genes10120973

Szklarczyk, D., Morris, J. H., Cook, H., Kuhn, M., Wyder, S., Simonovic, M., Santos, A., Doncheva, N. T., Roth, A., Bork, P., Jensen, L. J., & von Mering, C. (2017). The STRING database in 2017: quality-controlled protein–protein association networks, made broadly accessible. Nucleic Acids Research, 45 (D1), D362–D368. https://doi.org/10.1093/nar/gkw937

Thelen, J. J., & Ohlrogge, J. B. (2002). Metabolic Engineering of Fatty Acid Biosynthesis in Plants. Metabolic Engineering, 4(1), 12–21. https://doi.org/10.1006/mben.2001.0204

Van, K., & McHale, L. (2017). Meta-Analyses of QTLs Associated with Protein and Oil Contents and Compositions in Soybean [Glycine max (L.) Merr.] Seed. International Journal of Molecular Sciences, 18(6), 1180. https://doi.org/10.3390/ijms18061180

Vaughn, J. N., Nelson, R. L., Song, Q., Cregan, P. B., & Li, Z. (2014). The Genetic Architecture of Seed Composition in Soybean Is Refined by Genome-Wide Association Scans Across Multiple Populations. G3&#58; Genes|Genomes|Genetics, 4(11), 2283–2294. https://doi.org/10.1534/g3.114.013433

Voelker, T., & Kinney, A. J. (2001). Variations in the biosynthesis of seed-storage lipids. Annual Review of Plant Physiology and Plant Molecular Biology, 52(1), 335–361. https://doi.org/10.1146/annurev.arplant.52.1.335

Wang, T., Xing, J., Liu, X., Liu, Z., Yao, Y., Hu, Z., Peng, H., Xin, M., Zhou, D. X., Zhang, Y., & Ni, Z. (2016). Histone acetyltransferase general control non-repressed protein 5 (GCN5) affects the fatty acid composition of Arabidopsis thaliana seeds by acetylating fatty acid desaturase3 (FAD3). Plant Journal, 88(5), 794–808. https://doi.org/10.1111/tpj.13300

Wang, X., & Komatsu, S. (2017). Improvement of Soybean Products Through the Response Mechanism Analysis Using Proteomic Technique. In Advances in Food and Nutrition Research (1st ed., Vol. 82, pp. 117–148). Elsevier Inc. https://doi.org/10.1016/bs.afnr.2016.12.006

Wen, Z., Boyse, J. F., Song, Q., Cregan, P. B., & Wang, D. (2015). Genomic consequences of selection and genome-wide association mapping in soybean. BMC Genomics, 16(1), 671. https://doi.org/10.1186/s12864-015-1872-y

Xu, C., & He, C. (2007). The rice OsLOL2 gene encodes a zinc finger protein involved in rice growth and disease resistance. Molecular Genetics and Genomics, 278(1), 85–94. https://doi.org/10.1007/s00438-007-0232-2

Yang, S., Miao, L., He, J., Zhang, K., Li, Y., & Gai, J. (2019). Dynamic transcriptome changes related to oil accumulation in developing soybean seeds. International Journal of Molecular Sciences, 20 (9). https://doi.org/10.3390/ijms20092202

Yao, Y., You, Q., Duan, G., Ren, J., Chu, S., Zhao, J., Li, X., Zhou, X., & Jiao, Y. (2020). Quantitative trait loci analysis of seed oil content and composition of wild and cultivated soybean. BMC Plant Biology, 20(1), 1–13. https://doi.org/10.1186/s12870-019-2199-7

Zhang, D., Lü, H., Chu, S., Zhang, H., Zhang, H., Yang, Y., Li, H., & Yu, D. (2017). The genetic architecture of water-soluble protein content and its genetic relationship to total protein content in soybean. Scientific Reports, 7(1), 5053. https://doi.org/10.1038/s41598-017-04685-7

Zhang, D., Zhang, H., Hu, Z., Chu, S., Yu, K., Lv, L., Yang, Y., Zhang, X., Chen, X., Kan, G., Tang, Y., An, Y. C., & Yu, D. (2019). Artificial selection on GmOLEO1 contributes to the increase in seed oil during soybean domestication. PLOS Genetics, 15(7), e1008267. https://doi.org/10.1371/journal.pgen.1008267

Zhang, J., Wang, X., Lu, Y., Bhusal, S. J., Song, Q., Cregan, P. B., Yen, Y., Brown, M., & Jiang, G.-L. (2018). Genome-wide Scan for Seed Composition Provides Insights into Soybean Quality Improvement and the Impacts of Domestication and Breeding. Molecular Plant, 11(3), 460–472. https://doi.org/10.1016/j.molp.2017.12.016

Zhang, L., Wang, S. B., Li, Q. G., Song, J., Hao, Y. Q., Zhou, L., Zheng, H. Q., Dunwell, J. M., & Zhang, Y. M. (2016). An integrated bioinformatics analysis reveals divergent evolutionary pattern of oil biosynthesis in high- and low-oil plants. PLoS ONE, 11(5), 1–24. https://doi.org/10.1371/journal.pone.0154882

Zhang, T., Song, C., Song, L., Shang, Z., Yang, S., Zhang, D., Sun, W., Shen, Q., & Zhao, D. (2017). RNA sequencing and coexpression analysis reveal key genes involved in α-linolenic acid biosynthesis in Perilla frutescens seed. International Journal of Molecular Sciences, 18(11), 1–16. https://doi.org/10.3390/ijms18112433

Zhang, Y., He, J., Wang, H., Meng, S., Xing, G., Li, Y., Yang, S., Zhao, J., Zhao, T., & Gai, J. (2018). Detecting the QTL-Allele System of Seed Oil Traits Using Multi-Locus Genome-Wide Association Analysis for Population Characterization and Optimal Cross Prediction in Soybean. Frontiers in Plant Science, 9 (December). https://doi.org/10.3389/fpls.2018.01793

Zhang, Y. Q., Lu, X., Zhao, F. Y., Li, Q. T., Niu, S. L., Wei, W., Zhang, W. K., Ma, B., Chen, S. Y., & Zhang, J. S. (2016). Soybean GmDREBL Increases Lipid Content in Seeds of Transgenic Arabidopsis. Scientific Reports, 6(2), 1–13. https://doi.org/10.1038/srep34307

Zhao, X., Jiang, H., Feng, L., Qu, Y., Teng, W., Qiu, L., Zheng, H., Han, Y., & Li, W. (2019). Genome-wide association and transcriptional studies reveal novel genes for unsaturated fatty acid synthesis in a panel of soybean accessions. BMC Genomics, 20(1), 68. https://doi.org/10.1186/s12864-019-5449-z

Zhou, Z., Jiang, Y., Wang, Z., Gou, Z., Lyu, J., Li, W., Yu, Y., Shu, L., Zhao, Y., Ma, Y., Fang, C., Shen, Y., Liu, T., Li, C., Li, Q., Wu, M., Wang, M., Wu, Y., Dong, Y., … Tian, Z. (2015). Resequencing 302 wild and cultivated accessions identifies genes related to domestication and improvement in soybean. Nature Biotechnology, 33(4), 408–414. https://doi.org/10.1038/nbt.3096

Zombori, Z., Nagy, B., Mihály, R., Pauk, J., Cseri, A., Sass, L., Gábor, H. V., & Dudits, D. (2020). Ring-type e3 ubiqitin ligase barley genes (Hvyrg1–2) control characteristics of both vegetative organs and seeds as yield components. Plants, 9(12), 1–15. https://doi.org/10.3390/plants9121693

