## Supplementary figures for "Integrating omics approaches to discover and prioritize candidate genes involved in oil biosynthesis in soybean"

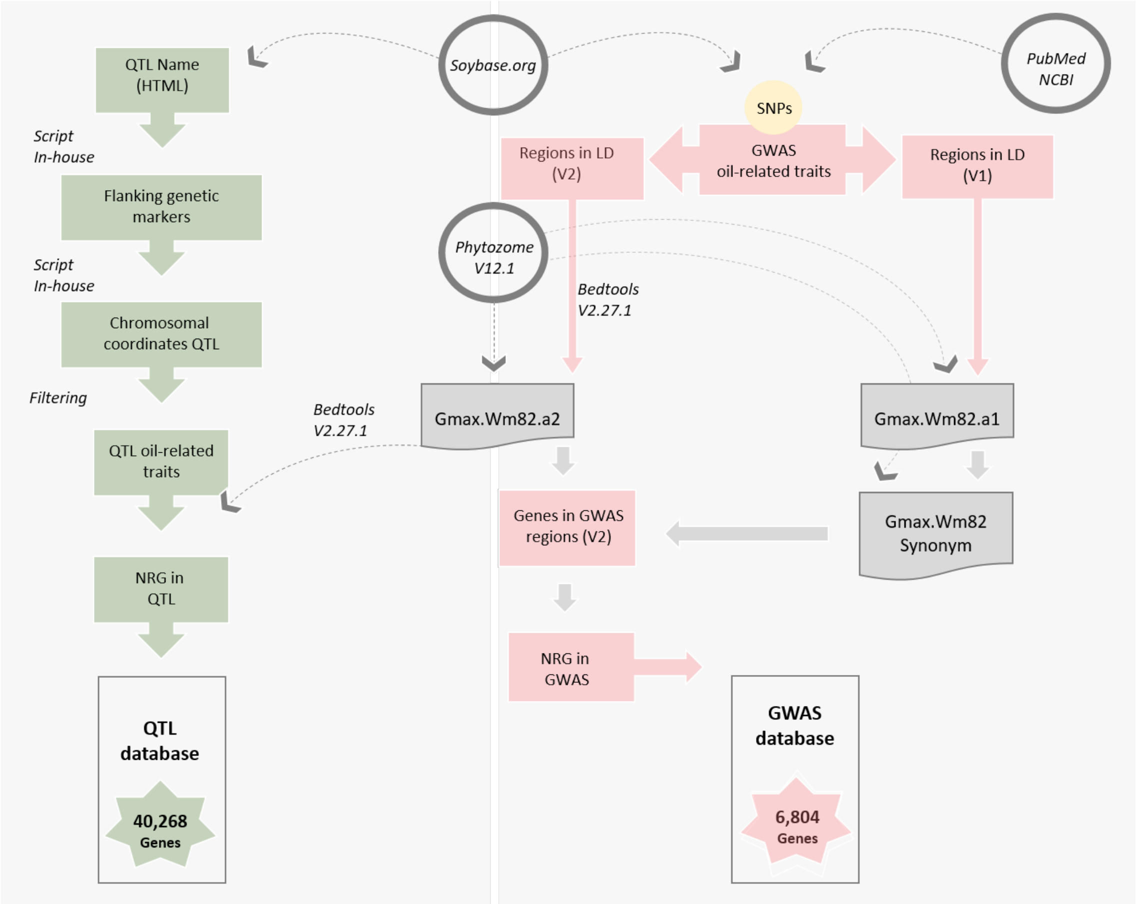


**Figure S1**. Workflow used to raise QTL and GWAS databases. **NRG**: Non-redundant genes; **LD**: linkage disequilibrium; **SNPs**: single nucleotide polymorphism.

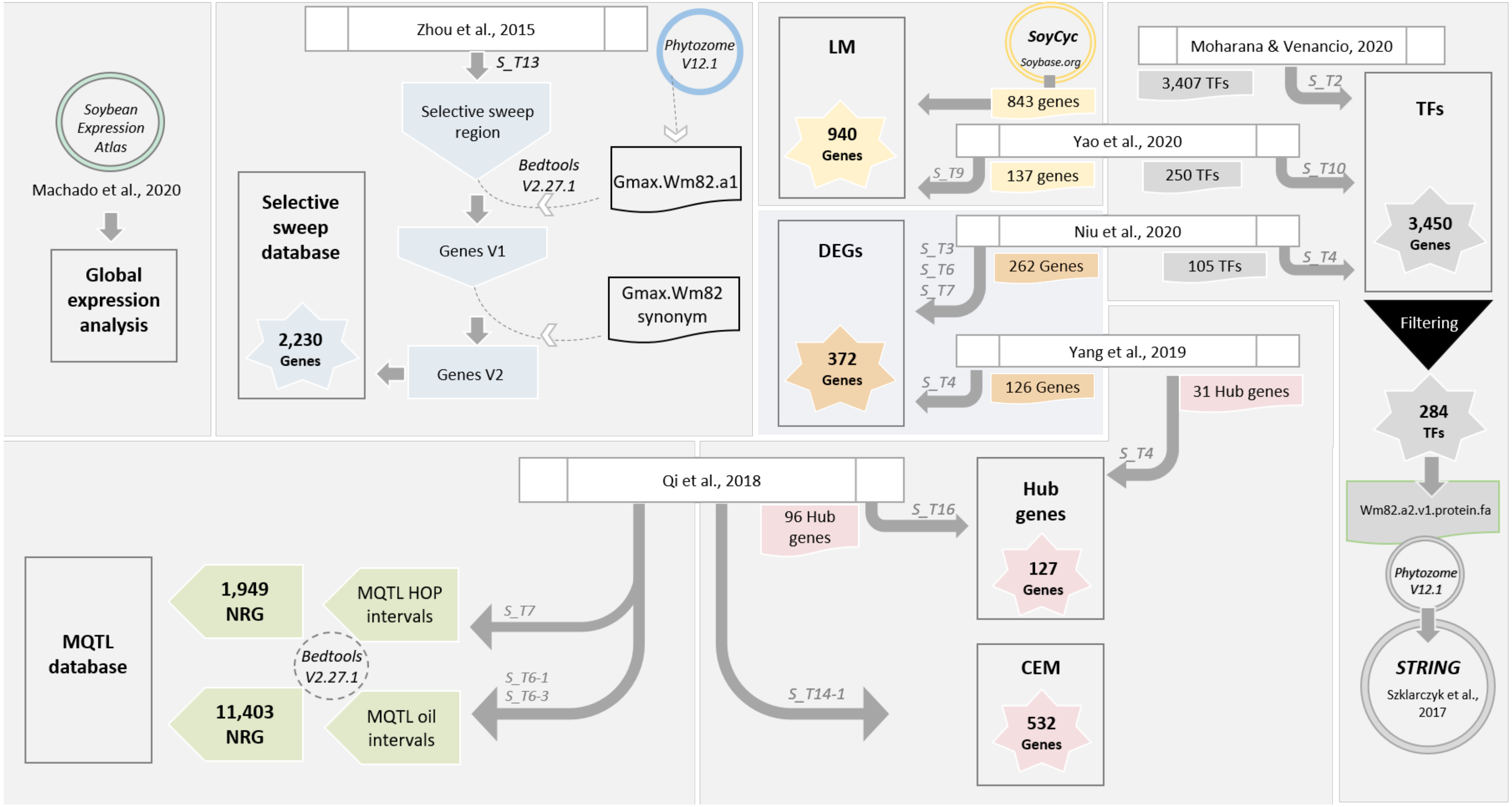


**Figure S2.** Workflow used to raise data that complemented the integrative analysis. **S_T*** means the supplementary tables which data were originally obtained; **LM**: lipid metabolism; **DEGs**: differentially expressed genes; **TFs**: transcription factors; **CEM**: coexpression modules; **HOP**: hotspots for oil and protein; **NRG**: non-redundant genes.

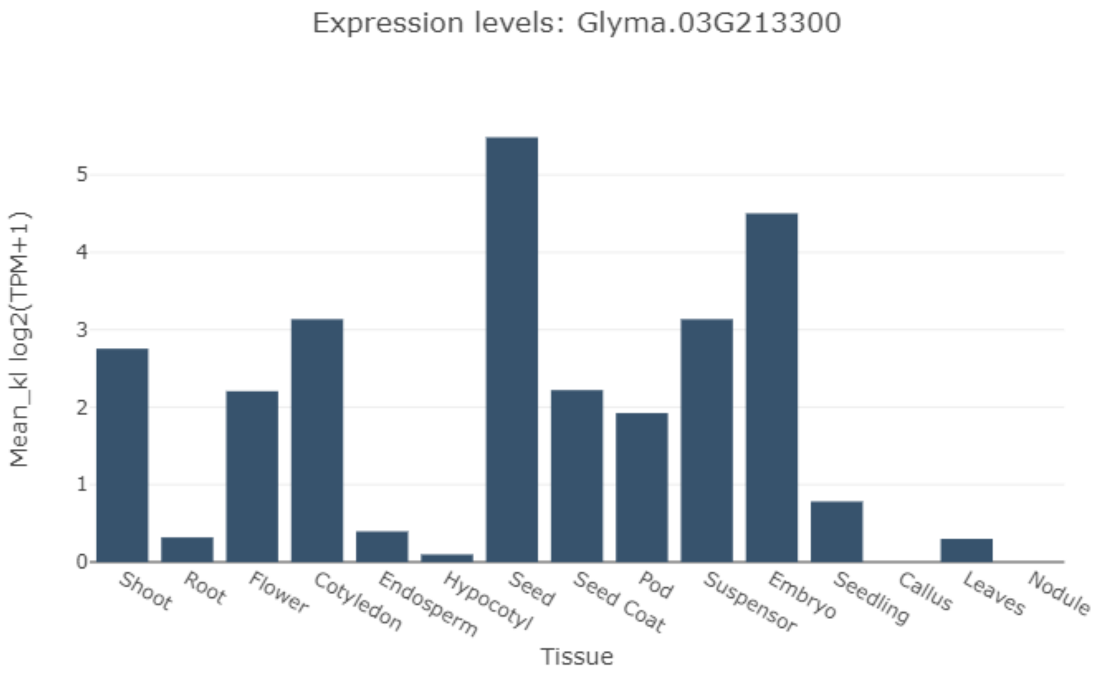


**Figure S3.** Average global expression of the membrane translocator protein - TSPO (Glyma.03G213300) using Kallisto as a method of estimating abundance of transcripts and normalization in transcripts per million (TPM). According to these data, the expression of TSPO is greater in soybean seed tissues. Source: http://venanciogroup.uenf.br/cgi-bin/gmax_atlas/index.cgi.

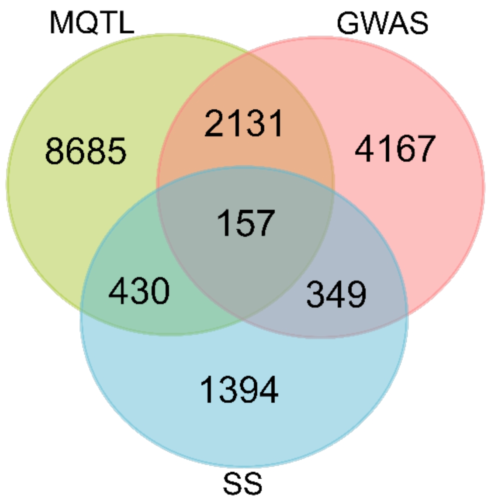


**Figure S4.** Venn diagram of genes in meta-QTL (**MQTL**), Genome Wide Association Studies (**GWAS**) and Selective Sweep regions (**SS**) database.

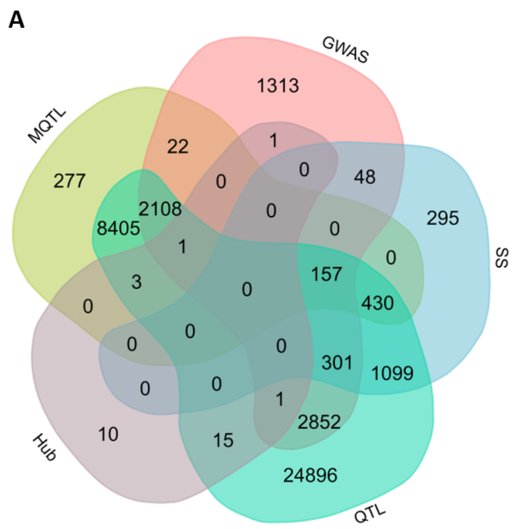


**Figure S5.** Venn diagram of hub genes in integrative analysis. **A.** Hub genes reported by Yang et al. 2019. Five of them are, at least, in two oil-related regions. **B.** Hub genes reported by Qi et al. 2018. Four of them are in MQTL, GWAS and selective sweep regions (**SS**).

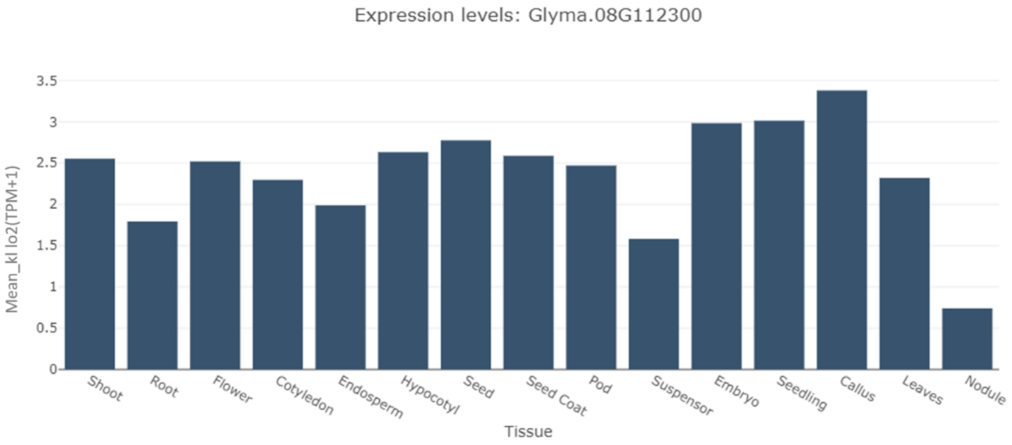


**Figure S6.** Average global expression of the Glyma.08G112300. As reported by Zhang et al. 2017, the expression of Glyma.08G112300 could be detected in different tissues. Thus, its role in seed protein composition needs further investigation. The estimation of transcript count was performed using kallisto (kl). Source: http://venanciogroup.uenf.br/cgi-bin/gmax_atlas/index.cgi.

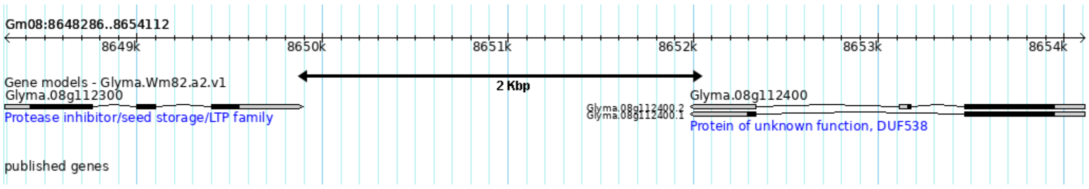


**Figure S7.** Position of Glyma.08G112400 in relation to Glyma.08G112300, both found in our investigations. Source: soybase.org
